# Repetitive DNA reeling by the Cascade-Cas3 complex in nucleotide unwinding steps

**DOI:** 10.1101/207886

**Authors:** Luuk Loeff, Stan J.J. Brouns, Chirlmin Joo

## Abstract

CRISPR-Cas loci provide an RNA-guided adaptive immune system against invading genetic elements. Interference in type I systems relies on the RNA-guided surveillance complex Cascade for target DNA recognition and the trans-acting Cas3 helicase/nuclease protein for target degradation. Even though the biochemistry of CRISPR interference has been largely covered, the biophysics of DNA unwinding and coupling of the helicase and nuclease domains of Cas3 remains elusive. Here we employed single-molecule FRET to probe the helicase activity with a high spatiotemporal resolution. We show that Cas3 remains tightly associated with the target-bound Cascade complex while reeling in the target DNA using a spring-loaded mechanism. This spring-loaded reeling occurs in distinct bursts of three base pairs, that each underlie three successive 1-nt unwinding events. Reeling is highly repetitive, compensating for inefficient nicking activity of the nuclease domain. Our study reveals that the discontinuous helicase properties of Cas3 and its tight interaction with Cascade ensure well controlled degradation of target DNA only.

## Introduction

Prokaryotes mediate defense against invading genetic elements using RNA guided adaptive immune systems that are encoded by CRISPR (clustered regularly interspaced short palindromic repeats)-Cas (CRISPR-associated) loci^1,2^. In the type I system, the most ubiquitous CRISPR-Cas system^3^, foreign DNA targets (called protospacers) are recognized by the CRISPR RNA (crRNA)-guided surveillance complex Cascade^4^. Recognition of double stranded DNA targets results in the formation of an R-loop, in which the crRNA hybridizes with the complementary target strand and the non-complementary strand of the DNA is displaced (nontarget strand)^5–10^. This R-loop formation triggers a conformational change in the Cascade complex^6,10–12^ and leads to the recruitment of the Cas3 protein for subsequent target degradation^13–15^.

The *E. coli* Cas3 protein consists of two domains: a N-terminal metal-dependent histidine-aspartate (HD) nuclease domain and a C-terminal superfamily 2 helicase domain^3,13,16–19^. Cas3 is activated by the Cascade-marked R-loop, where it cleaves the displaced nontarget strand ~11 nucleotides into the R-loop region^16,20^. Driven by ATP, Cas3 then moves along the nontarget strand in a 3’ to 5’ direction, while catalyzing cobalt-dependent DNA degradation^14,16,20,21^. Subsequently, Cas3 generates degradation products that are close to spacer length and enriched for PAM-like NTT sequences in their 3’ ends^22^. This makes a considerable fraction of the degradation products suitable substrates for integration by the Cas1-Cas2 integrases into the CRISPR locus^22,23^. Even though the biochemistry of CRISPR interference has been largely covered, the biophysics of DNA unwinding by Cas3 remains elusive. In particular, how the helicase domain tunes the property of nuclease HD domain for spacer integration and how this process takes place in concert with Cascade.

## Results

### Single-molecule observation of DNA reeling by Cas3

We set out to understand how Cas3 unwinds dsDNA substrates. To date, two models prevail for DNA unwinding by the Cas3 helicase: a translocation model and a reeling model. In the translocation model, Cas3 breaks its contacts with the Cascade complex while unwinding the DNA. Thereby Cas3 translocates away from the Cascade binding site and degrades single-stranded DNA fragments along the way (Fig 1a)^13–16,21,24^. In the reeling model, Cas3 and Cascade remain in tight contact while Cas3 unwinds the DNA, which may result in loops in the target strand (Fig 1a)^21,25^. To distinguish between these two models, we sought to visualize the DNA unwinding activity of Cas3 with a high spatiotemporal resolution.

**Figure 1.**
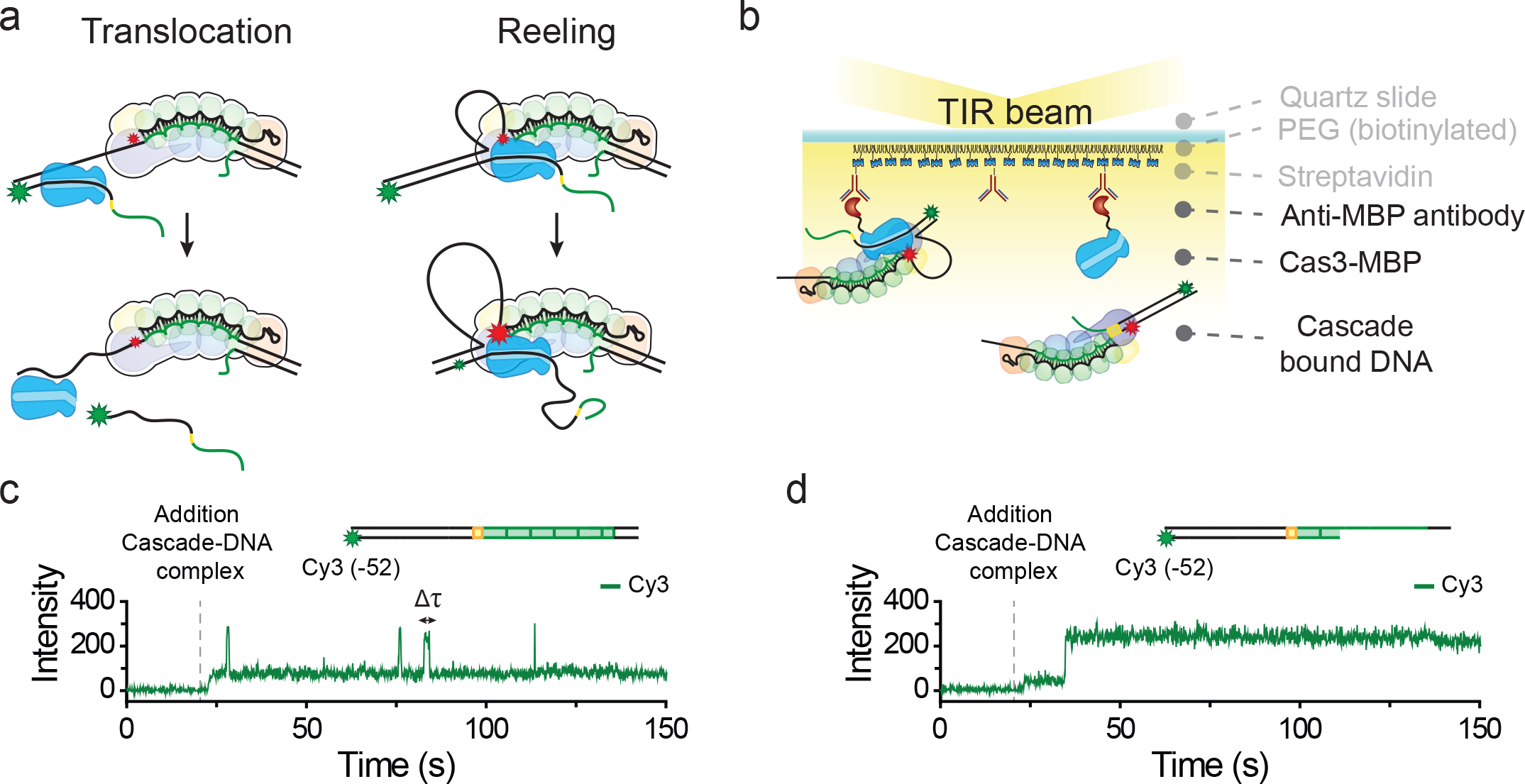
Single-molecule visualisation of the interaction between Cas3 and Cascade. **(a)** Schematic of two distinct model for DNA unwinding by Cas3. In the translocation model (left), Cas3 breaks its contacts with Cascade while it unwinds DNA. This results in translocation away from the Cascade target site. During DNA reeling (right), Cascade and Cas3 remain tightly associated while Cas3 pulls on the DNA. As a result, loops are formed in the target strand. The appearance of FRET during translocation or reeling is indicated by the size of the star: low FRET, large green star) or high FRET, large red star. **(b)**Schematic of a single-molecule FRET assay used to probe the interaction between Cas3 and Cascade. **(c)** A representative time trace of the initial interaction between Cas3 and Cascade bound to a cognate target labelled with a single fluorophore (Cy3). **(d)** A representative time trace of the initial interaction between Cas3 and Cascade bound to a partial duplex target labelled with a single fluorophore (Cy3).

To visualize DNA unwinding by Cas3, we developed an assay based on single-molecule Förster resonance energy transfer (smFRET). In brief, anti-maltose binding protein (MBP) antibodies were anchored to the surface of a polyethylene glycol (PEG)-coated slide through biotin-streptavidin linkage followed by tethering of MBP-fused Cas3 monomers (Fig. 1b & Extended Data Fig. 1a-c)^25^. Notably, the immobilization of Cas3 did not appreciably affect its capability to degrade dsDNA substrates (Extended Data Fig. 1c-g). Next, the antibody-tethered Cas3 molecules were presented to Cascade complexes bound to dye-labeled dsDNA substrates and their interactions were probed in real time using total internal reflection fluorescence (TIRF) microscopy (Fig. 1b). We first explored the interaction of Cas3 with Cascade complexes that were bound to a fully complementary dsDNA target. When the complexes were introduced in absence of cobalt, transient interactions were observed with a dwell-time (Δ*τ*) of 1.63 ± 0.236 s, which reflect the initial interaction between the Cse1 subunit of the Cascade complex and the Cas3 protein (Fig. 1c & Extended Data Fig. 2a)^15,25^. This finding is consistent with DNA curtain experiments where no stable interaction between Cascade and Cas3 was observed when cobalt was omitted from the assay^21,25^. To focus on the Cas3 helicase activity, experiments were performed in the absence of cobalt, unless stated otherwise.

When the same experiment was repeated but with a partial dsDNA construct that mimicked the nicked R-loop reaction intermediate formed by Cas3 (Fig 2a), a stable interaction was observed between Cascade and Cas3. This interaction lasted throughout the time course of the experiment and followed photo bleaching kinetics (Fig. 1d & Extended Data Fig. 2b). This suggest that the initial nick made by Cas3 facilitates loading of the helicase domain, which stabilizes the interaction between Cas3 and the Cascade complex. Notably, the appearance of fluorescence signals was not observed when Cascade was omitted from the assay, confirming that Cas3 exclusively interacts with DNA in a Cascade-dependent manner (Extended Data Fig. 2c & 2d)^8,14,15,26^.

**Figure 2.**
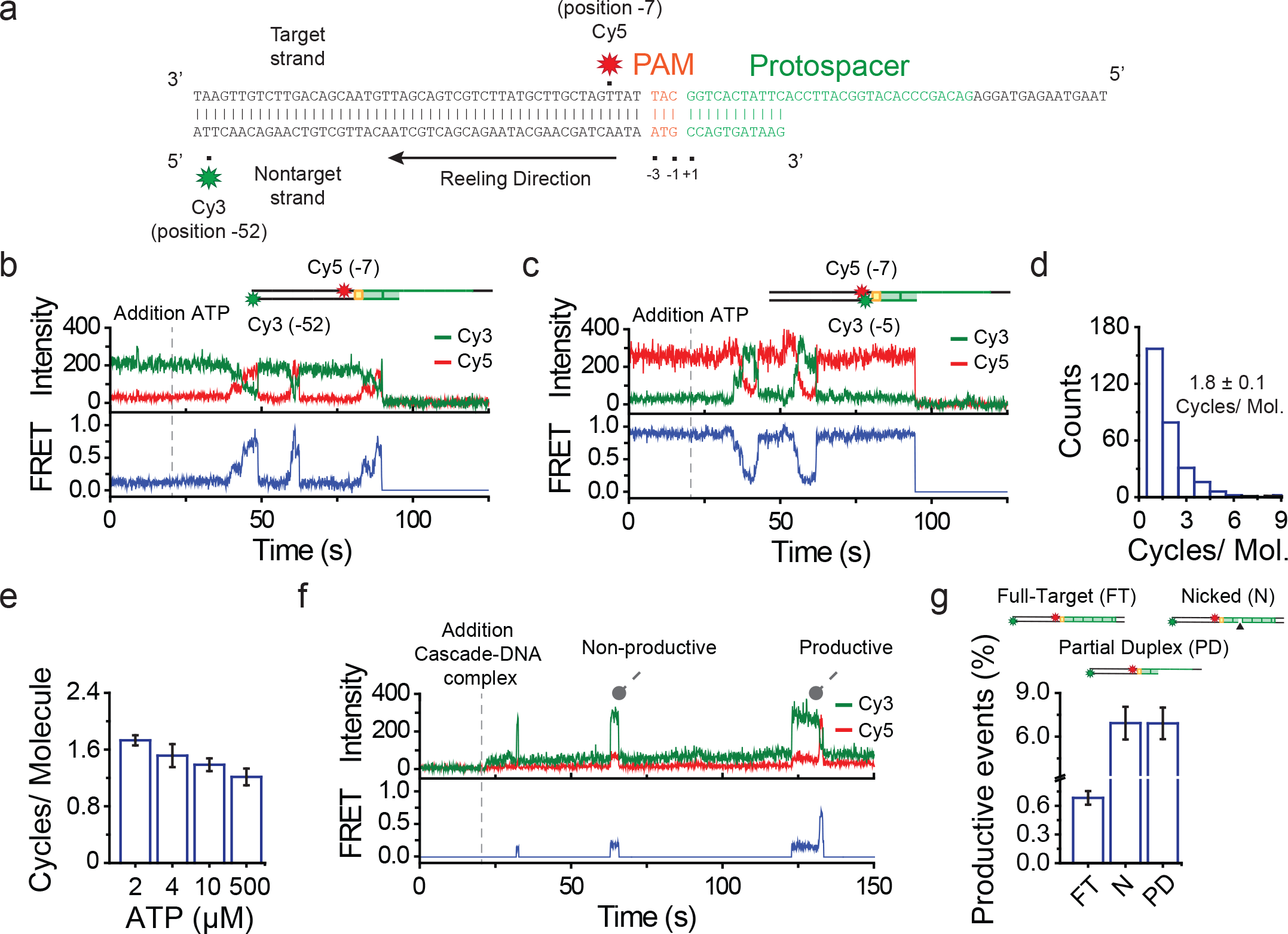
Real-time observation of DNA reeling by Cas3. **(a)** Partial duplex DNA constructs consist of a PAM (orange), protospacer (green) and two flanks of 50 nt and 15 nt (black). Cy5 (red star) was attached to position −7 of the target strand and Cy3 (green star) to position −52 of the nontarget strand. **(b)** A representativetime trace of donor (Cy3, green) and acceptor (Cy5, red) fluorescence and corresponding FRET (blue) exhibiting multiple reeling events. ATP (2 μM) was added at t = 20s (dashed gray line). **(c)** A representative time trace for a construct with Cy5 (red star) attached to position −7 of the target strand and Cy3 (green star) to position −5 of the nontarget strand. ATP (2 μM) was added at t = 20s (dashed gray line). **(d)** A histogram representing the number of reeling cycles for each molecule. Error represents the standard error of the mean (SEM) from three independent measurements (N=3). **(e)** Quantification of the number of reeling cycles per molecule at various ATP concentrations. Error bars represent the SEM (N=3). **(f)** Representative time traces of donor (Cy3, green) and acceptor (Cy5, red) fluorescence and corresponding FRET (blue) obtained by tracking the interactions by Cas3 and Cascade in real time. Cascade bound DNA, ATP (500 μM) and Co^2+^ (10 μM) were added at t = 20s. **(g)** Quantification of the number of productive binding events for three distinct DNA constructs. Cy5 (red star) was attached to position −7 of the target strand and Cy3 (green star) to position −52 of the nontarget strand. Black triangle indicates the position of the nick (11 nt away from PAM).

To synchronize the initiation of DNA unwinding, experiments were continued with the partial dsDNA construct. The DNA substrate was labelled with a donor (Cy3) and an acceptor (Cy5) dye that were positioned such that it could report on reeling along the non-target strand via an increase in FRET (Fig. 1a, Fig 2a & Extended Data Table 1). The fluorescent probes where conjugated to the DNA using an amino-C6-linker, which has been shown not to interfere with the translocation and unwinding by helicases^27–30^. The target strand was labeled with Cy5 at nucleotide −7, which position is fixed near the Cascade complex^8,10^. The Cy3 dye was positioned further upstream of the PAM at position −52 such that high FRET would be observed upon reeling along the non-target strand by Cas3 (Fig. 1a). In absence of ATP, no FRET was observed between the donor and acceptor fluorophore, resulting in FRET values that were indistinguishable from background signals (E = 0.18) (Fig. 1b & Extended data Fig. 2e).

Upon introduction of ATP into the microfluidic chamber, a large fraction of the Cas3 molecules (201 out of 438 molecules) showed a gradual increase in FRET, which is consistent with reeling along the non-target strand (Fig. 2b & Extended data Fig. 3a). For remaining molecules, FRET stayed within background levels (E = 0.18). We hypothesize that these molecules either failed to initiate unwinding within our observation time (3.5 min) or did not reel the DNA beyond the FRET range of approximately 20 base pairs (bp) (Extended data Fig. 8a & 8b). Consistent with the second hypothesis, the probability of reeling scaled exponentially with the distance to the target-site (Fig. 3d). This data shows that Cas3 remains anchored to Cascade while reeling the DNA. Notably, translocation of Cas3 away from the Cascade target site^13–16,21,24^ would have been manifested by a rapid loss of the total fluorescence signal (Fig. 1a), which was not observed under our tension free experimental conditions (Extended Data Fig. 2b). This is supported by the finding of tension dependent Cas3-Cascade rupture^25^.

**Figure 3.**
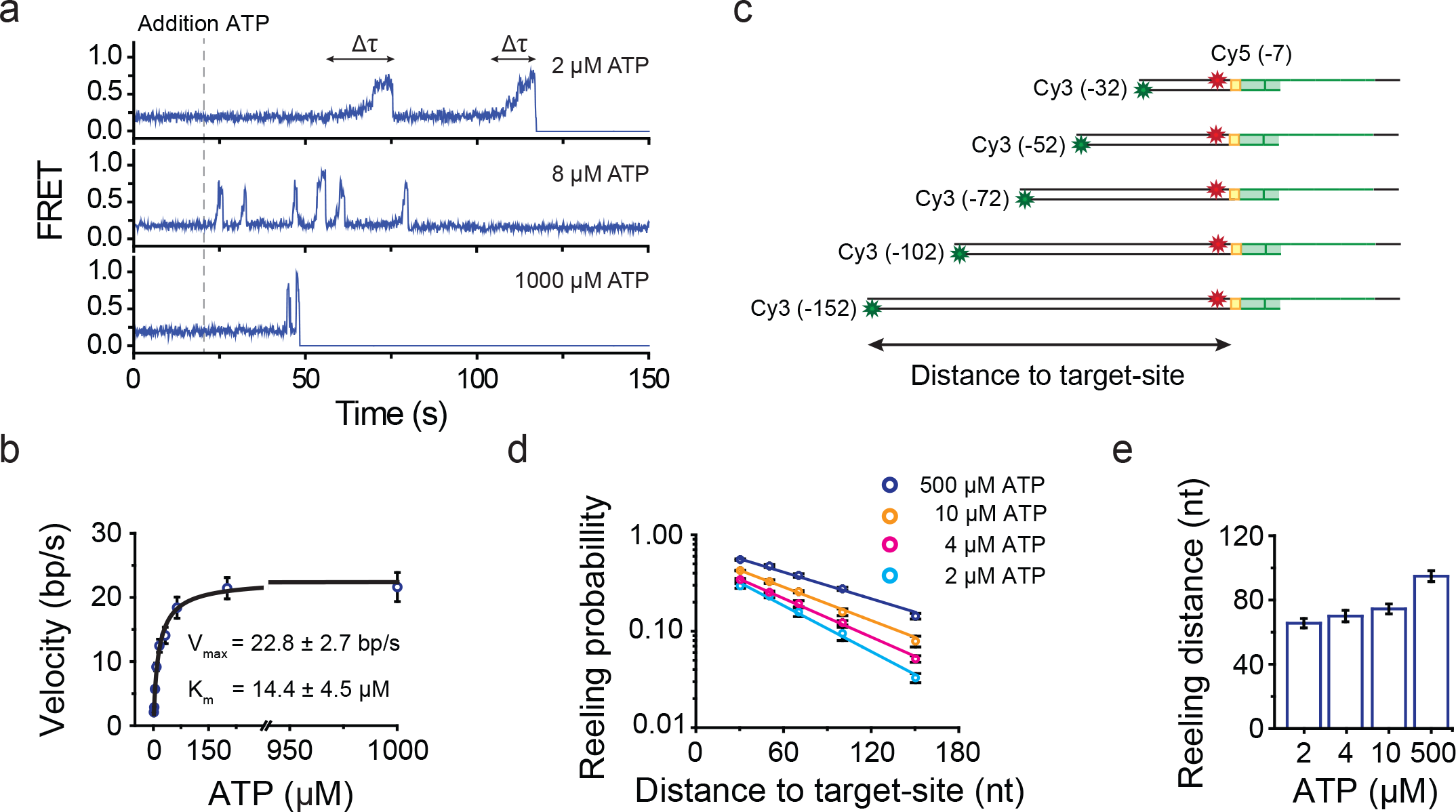
Velocity and processivity of the Cas3 helicase. **(a)** Representative FRET traces obtained at various ATP concentrations. **(b)** Michaelis-Menten fit (black line) of the velocity (bp/s) plotted against the ATP concentration (1, 2, 4, 8, 16, 32, 64, 200 and 1000 μM ATP). Error bars represent the 95% confidence interval obtained through bootstrap analysis. **(c)** Schematic overview of constructs used to determine the mean reeling distance. Cy5 (red star) was attached to position −7 of the target strand and Cy3 (green star) was positioned at the end of the nontarget strand. **(d)** Reeling probability over the distance to target-site at various ATP concentrations. Error bars represent the SEM (N=3). Solid lines represent a single-exponential fit used to determine the mean reeling distance. **(e)** Bar plot with average reeling distance (nt) at various ATP concentrations. Error bars represent the SEM (N=3).

To confirm DNA reeling along the non-target strand, we tested various alternative immobilization and labelling schemes. When the DNA (Extended data Fig. 3c & 3d) or Cascade (Extended data Fig. 3e & 3f) was immobilized or when the donor and acceptor dyes were swapped (Extended Data Fig. 4a, d), identical behavior was observed. Next, an alternative labelling scheme was tested, with a donor and acceptor dye at position −5 of the nontarget strand and −7 of the target strand, respectively (Extended Data Fig. 2f), was tested. This construct initially yielded high FRET (E = 0.8, Extended Data Fig. 2f) and should lead to a decrease in FRET when reeling is triggered. In agreement with our expectation, FRET decreased upon introduction of ATP (Fig 2c & Extended Data Fig. 3b). The same observation was made using PIFE (protein-induced fluorescence enhancement), albeit with lower resolution (Extended Data Fig. 3g). In contrast, when a construct was used that was designed to detect DNA reeling along the target strand (Cy3 position −52 target strand and Cy5 position −7 nontarget strand, Extended Data Fig. 4b) or when the PAM proximal and PAM distal flank were swapped (Extended Data Fig. 4c), a change in FRET was not observed (Extended Data Fig. 4e & 4f). These control experiments support the model that Cas3 remains anchored to Cascade^25^ when reeling in the 3’ end of nontarget strand, resulting in DNA loops in the target strand.

Our real-time analysis of DNA reeling by Cas3 revealed, that Cas3 could go through multiple cycles of reeling on a single substrate, by slipping back to its initial location (Fig. 2b & 2c)^25^. Analysis of this repetitive behavior showed that Cas3 undergoes an average of 1.8 ± 0.1 cycles per substrate (Fig. 2d). Interestingly, the number of reeling cycles per molecule decreased with an increase of ATP, reaching average unwinding frequency of 1.2 ± 0.1 cycles per molecule at saturating levels of ATP (Fig. 2e). This data suggests that Cas3 is more effective in displacing the nontarget strand away from the Cas3-Cascade complex at higher levels of ATP, which is likely a result of using short DNA oligo’s. Consistent with this hypothesis, the dwell time of the looping population that reached the end of the substrate was ~3 times shorter as compared to the seemingly static population (Extended Data Fig. 2b).

### Cas3 exhibits sparse nuclease activity

Next, we sought to how the repetitive reeling behavior is coordinated with the nuclease activity of the HD domain. Previous bulk experiments have shown that Cas3 degrades the nontarget strand while it moves along the DNA^16,20^. Therefore, we hypothesized that activation of the nuclease domain, by the addition of Co^2+^ would result in a stark decrease in the number of reeling cycles per molecule. However, no change in the number of cycles per molecule was observed when the nuclease domain was activated (Extended Data Fig. 3h), indicating that little nicking had occurred. Moreover, the addition of free Cas3 into the assay did not alter the behavior of Cas3 (data not shown). Consistent with this finding marginal degradation of the nontarget strand was observed in a biochemical assay on DNA oligos (Extended Data Fig. 1h).

To obtain a more quantitative estimate on the cleavage activity of Cas3, the initial interaction between Cas3 and Cascade was probed (Fig 2f). When Cascade bound to a full target substrate, without the initial nick, was introduced, only 0.7 ± 0.1% of the binding events resulted in DNA reeling (Fig 2f & 2g). However, when Cascade was introduced bound to a substrate mimicking the nicked intermediate, the number of reeling events increased with an order of magnitude (6.9 ± 1.1%, Fig 2f & 2g). This suggest that the HD nuclease domain intrinsically exhibits a sparse nuclease activity, which contrasts previously published bulk data^14–16,20,22,31^. Those bulk measurements were performed using a 10- to 500-fold excess of Cas3^14–16,20,22,31^ that facilitated initial nicking and loading of the Cas3 helicase, whereas here we used Cas3 in nano-molar concentrations. Our findings imply that the Cas3 protein compensates for the sparse nuclease activity by repeatedly reeling ssDNA substrates into the HD nuclease domain, which ensures DNA cleavage.

### Dynamics of DNA loop formation by Cas3

Next, we explored the molecular dynamics of DNA reeling by Cas3. To determine the unwinding rate of Cas3, we performed DNA unwinding assays at various ATP concentrations (Extended Data Fig. 5d-f). For every ATP concentration, the dwell time (Δ*τ*) of each unwinding event was extracted (Fig. 3a), followed by fitting of the histograms with a gamma distribution (Extended Data Fig. 5a-c). Consistent with other helicases^27,29,32^, the effective rate (k_effective_, 1/Δ*τ*) increased with increasing amounts of ATP, indicating that the reeling velocity increases with ATP (Fig 3b). When ATP was replaced with a non-hydrolysable ATP analog ATP-*γ*-S, the reeling activity of Cas3 was completely abrogated (Data Fig. 6a). To estimate the maximum velocity (V_max_) of Cas3, the effective rate was converted to apparent velocity in base pairs per second (bp/s, see Methods). By plotting the velocity over the ATP concentration and fitting the data with a Michaelis-Menten fit (Fig. 3b), a V_max_ = 22.8 ± 2.7 bp/s and K_m_ = 14.4 ± 4.5 μM was obtained. Notably, only a marginal change in velocity was observed when the nuclease domain was activated (Extended Data Fig. 5g), suggesting that the reeling activity of the helicase domain dominates over the DNA degradation by the nuclease domain.

Recent DNA curtain experiments suggested that Cas3 is a highly processive molecular motor^21^. However, given that the Cas3 nuclease exhibits sparse activity (Fig 2f & 2g), a highly processive motor would lead to single-stranded fragments that are much longer than the previously reported fragment size that is smaller than 200 nucleotides^16,22^. Therefore, we sought to determine the average reeling distance of Cas3 at saturating concentrations of ATP. To estimate the reeling distance, a series of DNA substrates were used with an increasing length of the PAM proximal flank, while moving the donor dye towards the end of each substrate (Fig. 3c). This set of constructs allowed for the determination of the probability that a Cas3 molecule reached the end of a DNA substrate within the observation time of 3.5 min.

Upon introduction of ATP, each construct yielded traces with identical behavior (Extended Data Fig. 7). However, we observed a decrease in the number of events with an increase in the flank length, suggesting that the reeling probability decreased (Fig 3d). When the length of the flank was increased to 150 nt, the reeling probability decreased to 0. 13 ± 0.1 (Fig 3d), suggesting that the majority of molecules formed loops smaller than 150 nt. To estimate the average unwinding distance, the reeling probability was plotted over the ATP concentration, followed by fitting each data series with a single-exponential decay. This yielded an average reeling distance of 95 ± 3 nt at a saturating ATP concentration (Fig. 3e). A decrease in the average reeling distance was observed when the ATP concentration was lowered (Fig. 3e). Notably, the addition of SSB did not alter the processivity of Cas3 (data not shown)^25^, implying that Cas3 may shelter the looped target strand. These observations are in good agreement previously reported bulk biochemical data, which showed that Cas3 generates degradation products in the range of 30 to 150 nucleotides that become smaller at low ATP concentrations^16,22,33^. Taken together, these results suggest that the helicase domain of Cas3 limits the fragment size by repeatedly generating a distribution ssDNA fragments with an average size of ~90 nt.

### Cas3 unwinds DNA in uniform steps

To understand what feature of the Cas3 helicase limits the reeling distance, we sought to understand the molecular mechanism by which Cas3 unwinds the DNA. Close inspection of the FRET events revealed that FRET increased with a distinct pattern, marked by plateaus at specific FRET levels (Fig. 4a). To elucidate this behavior, we employed an automated step-finder algorithm^34^ (Loeff et al., Method article in preparation) that yielded the average FRET value for each plateau and the size of each step in between the plateaus (Fig. 4a). Analysis of the average the FRET value for each plateau, resulted in a histogram with four distinct peaks that were evenly separated (ΔE=0.15) (Fig. 4b). To correlate these FRET values to distance in bp, we designed a series of DNA constructs, in which the distance between the dyes was systematically decreased (Extended Data Fig. 8a & 8b). The calibration experiment yielded a conversion factor, in which 1-bp corresponds to a ΔE=0.05 FRET change. Notably, this conversion factor is in line with previously published work^35^. Conversion of the FRET values suggests that Cas3 reels the DNA with regular 3-bp steps.

**Figure 4.**
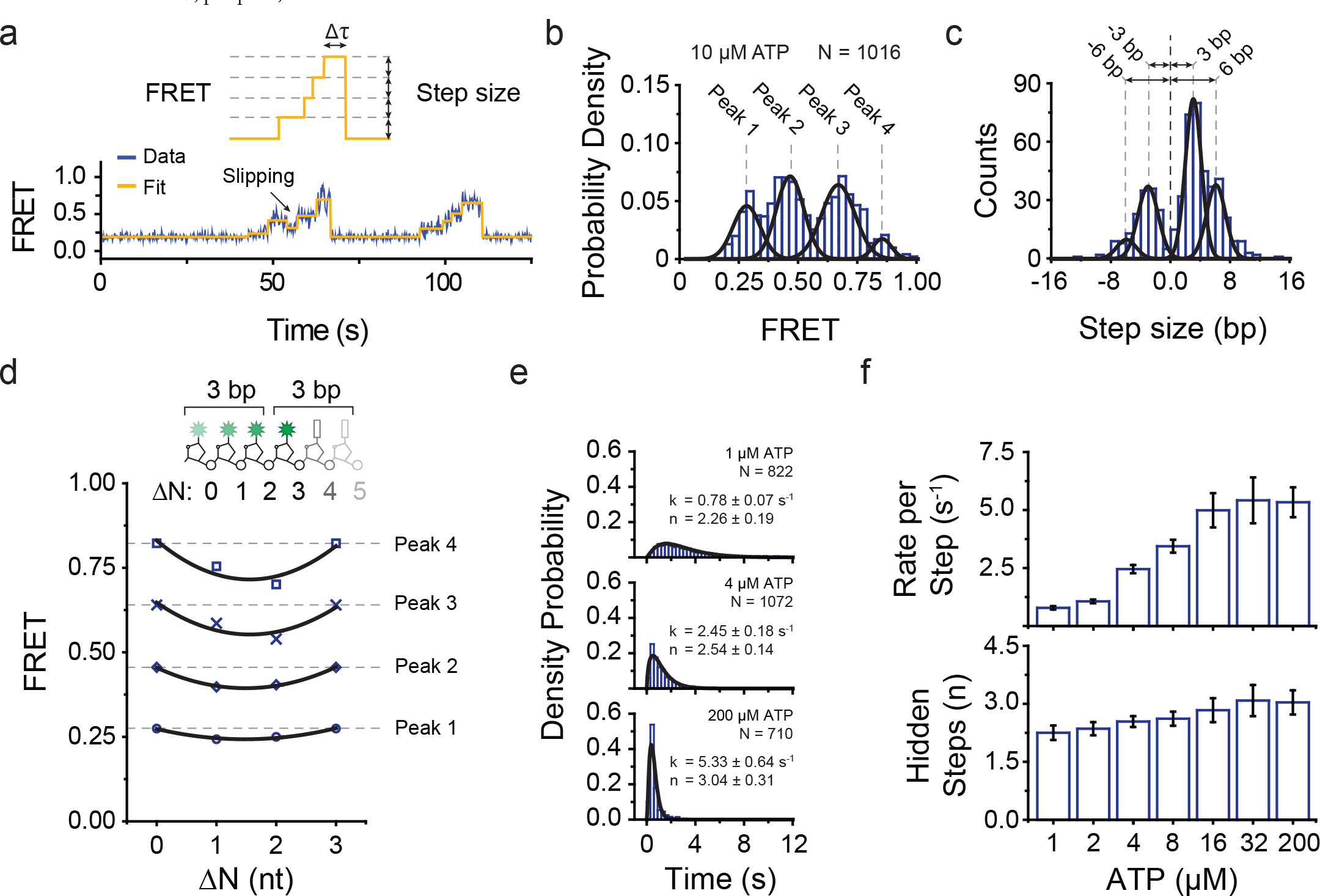
Cas3 reels DNA in uniform steps. **(a)** Representative FRET trace (dark blue) fitted with a step-finder algorithm (orange). **(b)** Distribution of FRET levels obtained through the step-finder algorithm. Black lines represent a Gaussian fit. (c) Distribution of step-sizes obtained through a step-finder algorithm. Black lines represent a Gaussian fit. Dashed grey lines indicate the centre of each peak. Positive values represent processive reeling whereas negative values represent slipping. **(d)** Location of the FRET levels for various positions of the donor dye. Given the remarkable regularity in the reeling pattern of Cas3, we hypothesized that when moving the donor dye from its original position (ΔN=0) by one or two nucleotides (ΔN=1 & ΔN=2, respectively) would shift the position of the observed plateaus at specific FRET levels. In contrast, moving the donor dyes by three nucleotides (ΔN=3) locates the dye at a similar position as ΔN=0 (inset) and should yield identical FRET levels. Consistent with our hypothesis, the constructs with a donor dye at position ΔN=1 & ΔN=2 shifted the peak positions towards lower FRET values, whereas the construct with a dye at position ΔN=3 yielded identical FRET levels as ΔN=0 (Extended Data Fig. 8f). **(e)** Dwell-time distributions of the FRET levels at various ATP concentrations. Data was fitted with a gamma distribution (solid line) to obtain the number of hidden steps (n) and rate (k). Error represents the 95% confidence interval obtained through bootstrap analysis. **(f)** Bar plots representing the number of hidden steps (n) and rate (k) that was obtained through fitting dwell-time histograms with a gamma distribution. Error represents the 95% confidence interval obtained through bootstrap analysis.

To further characterize the stepping behavior of Cas3, a histogram was plotted with the distribution of step-sizes, the distance between each plateau. The distribution of the step-sizes exhibited a major peak centered at a step-size of approximately 3-bp (Fig. 4c & Extended Data Fig. 8a-c), which is consistent with the histogram of average the FRET value for each plateau (Fig. 4b). Apart from the major peak at 3-bp, minor peaks that represented a multiplicity of this step-size (e.g. 6-bp) were observed (Fig. 4c), which became more prominent when the ATP concentration was increased (Extended Data Fig. 8d). These larger steps are likely a result of a series of events that occur faster than our time resolution. Consistent with this hypothesis, a histogram of the average FRET levels at saturating concentration of ATP was skewed towards the high FRET states (Extended Data Fig. 8e). We confirmed the 3-bp step by designing a set of constructs in which the donor dye was shifted by 1, 2, or 3 nt (ΔN=1, ΔN=2 & ΔN=3) from the standard construct (ΔN=0), resulting in 3-nt periodicity in FRET histograms (Fig. 4d). These experiments provide strong evidence that Cas3 reels the DNA in distinct steps of 3-bp at a time.

Apart from steps that led to an increase in FRET, we also observed slipping events where the FRET signal abruptly dropped to intermediate levels (Fig. 4a). These events were represented as a negative value in our step-size analysis and showed a major peak centered at −3-bp (Fig. 4c). Besides the slipping events to intermediate levels, we also observed slipping events that returned to their initial FRET state (Fig. 2b & 2c). We speculate that these slipping events occur through miscoordination of the RecA-like domains of the Cas3 helicase^18,19^, leaving the DNA to zip back over a short or long distances. The short and long-range slipping events result in discontinuous and burst-like unwinding behavior, that allows Cas3 to repeatedly feed ssDNA fragments into the nuclease domain for further processing.

Finally, we questioned if the observed 3-bp steps would correspond to the elementary step-size of the Cas3 helicase. If Cas3 would unwind 3-bp upon the hydrolysis of a single ATP molecule, the dwell time (Δ*τ*, Fig. 4a) histogram of the FRET levels would follow a single-exponential decay. However, a dwell time histogram of the FRET levels showed non-exponential behavior and followed a gamma distribution (Fig. 4e). A fit of the histogram yielded a statistical description of the number of underlying hidden steps (n) and the rate per step (k) (Fig. 4e). At various ATP concentrations, we obtained n values that remained close to three hidden steps, whereas the rate increased with an increase of ATP (Fig. 4e, 4f & Extended Data Fig. 9). This analysis shows that each 3-bp step is composed of three hidden steps of 1-nt, suggesting that the elementary step-size of Cas3 is 1-nt. From this analysis, the model emerges that Cas3 successively unwinds three base pairs in 1-nt steps, using its RecA-like domains^18,19^. During these successive 1-nt translocation events, the DNA is held in place by the Cas3 protein, resulting in an abrupt 3-bp spring-loaded burst upon release (Supplemental movie 1).

## Discussion

CRISPR interference in the type I systems, relies on the interplay of multiple proteins to convey resistance against invading mobile genetic elements. Based on our results we propose a model for CRISPR interference, where the transacting Cas3 helicase/nuclease remains tightly anchored to the Cascade effector complex while reeling in the invader DNA (Fig 5). Together with the sparse nuclease activity of Cas3, this anchoring mechanism acts as a fail-safe to prevent the genome damaging effects of off-target DNA cleavage. Our data suggests that, Cas3 unwinding the DNA with its RecA-like domains, breaking open the dsDNA helix 1-nt at a time (Fig 5, inset & Supplemental movie 1). While unwinding the DNA in 1-nt steps, the DNA is held in place by the Cas3 protein until three of such steps have taken place. Unwinding of the third nucleotide triggers the release of the DNA, resulting in a spring-loaded burst that moves the helicase by three base pairs (Supplemental movie 1). Such spring-loaded unwinding has been observed for both helicases with a RecA-like fold (e.g. NS3^28^) and nucleases^36,37^ and presumably reflects a general feature of Cas3 proteins^17^. Finally, our data suggests that Cas3 limits its reeling distance through slipping (Supplemental movie 2), which allows for repetitive reeling that compensates for inefficient nicking activity of the nuclease domain. Furthermore, the tight interaction between Cascade and Cas3 at the onset of CRISPR-interference provides a fail-safe for detrimental off-target DNA cleavage.

**Figure 5.**
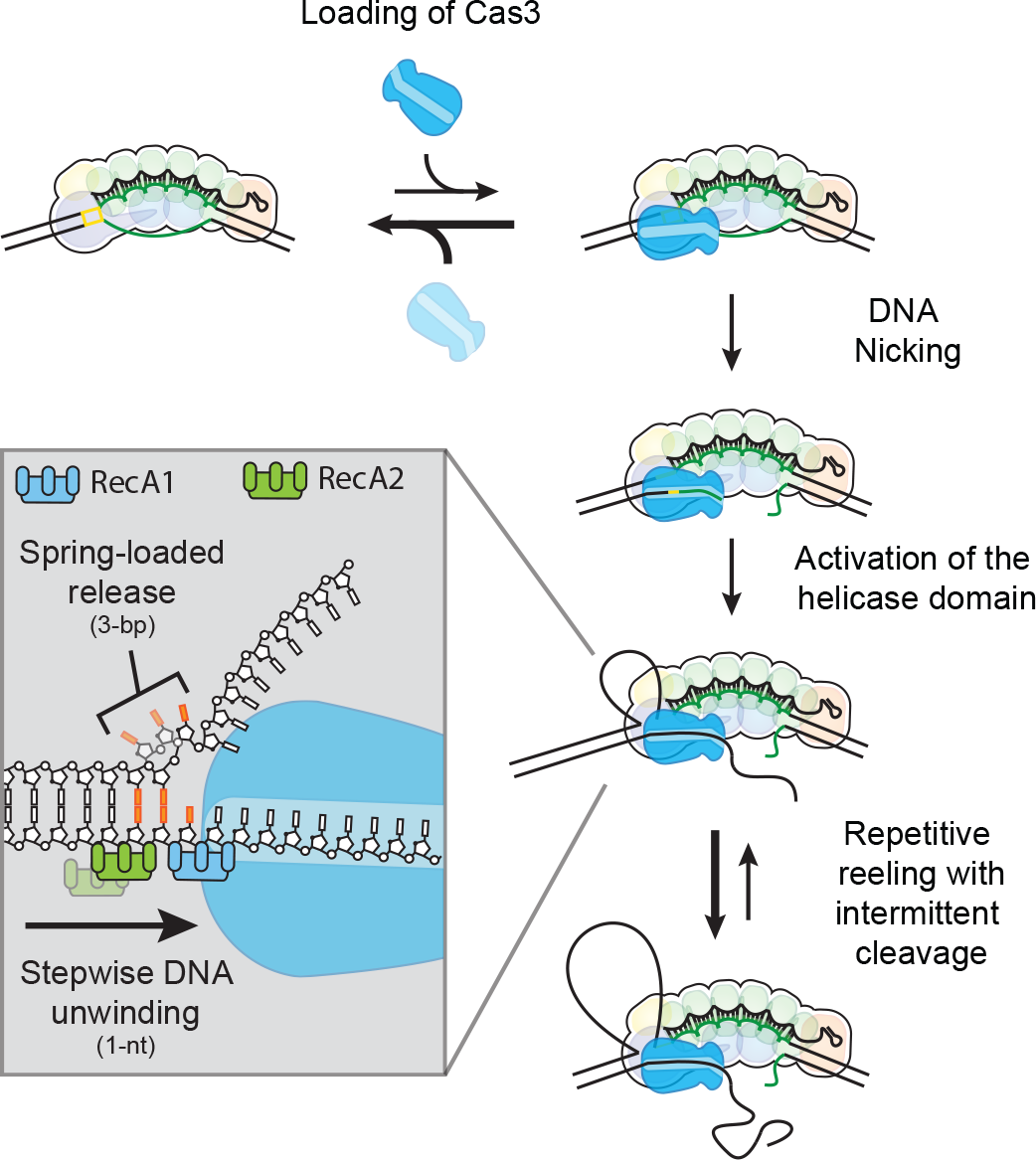
Model for CRISPR interference mediated DNA unwinding by Cas3. Model for CRISPR interference by Cas3. Interference starts with loading of the Cas3 protein onto the Cascade-DNA complex. Given the inherently sparse nuclease activity of Cas3, this may require multiple docking events. Once Cas3 nicks the R-loop, the loading of the helicase domain is facilitated that allows Cas3 and Cascade to form a stable complex. Upon hydrolysis of ATP, Cas3 initiates DNA reeling that takes place in distinct spring-loaded steps of 3-bp and underlies an elementary step size of 1-nt. Cas3 repeatedly feeds ssDNA into its nuclease domain, which generates a distribution of degradation products with an average size of ~90 nt.

## Acknowledgements

C.J. was funded by the Open Program of the Division for Earth and Life Sciences (822.02.008) and Vidi (864.14.002) of the Netherlands Organization for Scientific. S. J. J. B was funded by an LS6 ERC starting grant (639707) and a NWO VIDI grant (864.11.005). We would like to thank S. Bailey (Johns Hopkins University) for providing the Cas3 overexpression construct; We thank T. Blosser, T. Künne, M. Klein and R. McKenzie for critically reading this manuscript and we thank A.C. Haagsma, M. Klein, M. Depken, M. Szczepaniak and J. Kerssemakers for technical support.

## Author Contributions

L.L., S.J.J.B. and C.J. conceived the study; L.L performed experiments; L.L. analyzed data; L.L., S.J.J.B. and C.J. discussed the data and wrote the paper.

## Methods

### Protein preparation

Cascade was expressed in *E. coli* BL21 (DE3) and purified using strep-tag affinity chromatography, as described previously^5^. Purified Cascade complexes were aliquoted and flash frozen in liquid nitrogen for long-term storage at −80 °C. The nuclease-helicase Cas3 was produced and purified as described previously^16^ with the following modifications. BL21-AI cells were used for over-expression, and protein expression was induced with 0.5 mM IPTG and 0.2% L-Arabinose. The purification process was stopped after size exclusion chromatography and before the proteolytic removal of the Maltose Binding Protein (MBP) using the Tobacco Edge Virus protease^15^. MBP-Cas3 was aliquoted and flash frozen in liquid nitrogen before storage at −80 °C.

### Cas3 degradation Assays

After purification, Cas3 nuclease activity was initially tested by a non-specific degradation assay on M13mp8 singlestranded circular DNA (Extended Data Fig 1b). Non-specific nuclease activity was stimulated using Ni^+2^ ions as described previously^16^. To test specific degradation plasmid-based assays were performed in the presence of Cascade, ATP, Mg^+2^ and Co^+2^ ions (Extended Data Fig 1c-e), described previously by^16,22^. Similar conditions were used for oligo based degradation assays (Extended Data Fig 1h). In brief, 5 nM DNA was incubated with 50 nM Cascade and 100 nM Cas3 in buffer R (+10 μM CoCl_2_ and 2 mM ATP) for 30 minutes at 37°C. Samples were immediately quenched by adding stop solution (20 mM Tris-HCl pH 8.0, 2% SDS, 50 mM EDTA), after which protein was removed by incubating the samples for 1 hour with 10 μg/ ml proteinase K (Sigma) at 50°C. Subsequently, DNA was precipitated with ethanol and and loaded on 10% denaturing PAGE gels (8M urea) with formamide. Gels were run for 2.5 hour at 350 V, followed by imaging with the Typhoon trio (GE healthcare).

### DNA preparation

All the target dsDNA substrates that we used were bearing a protospacer, PAM, and two flanks of 50 and 15 nt (Fig. 2a, Extended Data Table 1). These synthetic DNA targets (Ella Biotech GmbH) were internally labelled with a monoreactive acceptor dye (Cy5, GE Healthcare) at dT-C6 on the target strand (complementary to the crRNA) and a monoreactive donor dye (Cy3, GE Healthcare) at dT-C6 on the nontarget strand (Fig 2a). After labelling, the ssDNA strands were annealed using a thermocycler (Biorad). To determine the initial FRET values of these constructs (Extended Data Fig. 2e-f & 4a-c), Cascade bound DNA was docked on the surface immobilized Cas3 molecules in absence of ATP.

### Single-molecule fluorescence data acquisition

The fluorescent label Cy3 and Cy5 were imaged using prism-type total internal reflection microscopy as described previously^6^ with slight modifications. After assembly of a microfluidic flow chamber, slides were incubated for 10 minutes with 5% Tween20 to further improve slide quality^38^ Next, the chamber was incubated with 20 μL streptavidin (0.1 mg/ml, S-888, Invitrogen) for 5 minutes followed by a washing step with 100 μL of buffer R. Anti-Maltose Binding Protein (anti-MBP) antibodies (M2155-09P, US biological life sciences) were specifically immobilized through biotin-streptavidin linkage by incubating the chamber with 40 μL of 10 μg/ ml anti-MBP antibodies for 5 minutes. Remaining unbound anti-MBP antibodies were flushed away with 100 μL buffer R. Subsequently, 100 μL of 10 nM Cas3-MBP was incubated in the chamber, allowing the Cas3-MBP molecules to bind the surface immobilized anti-MBP antibodies. After 5 minutes of incubation, unbound Cas3-MBP molecules were flushed away with 100 μL buffer R^imaging^ (50 mM HEPES (pH 7.5), 60 mM KCl, 10 mM MgCl_2_, 0.1 mg/mL glucose oxidase (G2133, Sigma), 4 μg/ml Catalase (10106810001, Roche) and 1 mM Trolox (((±)-6-Hydroxy-2,5,7,8-te-tramethylchromane-2-carboxylic acid, 238813, Sigma).

Cascade was incubated with 5 nM labelled dsDNA substrate with 50 nM Cascade for 5 minutes at 37°C. For docking experiments, pre-bound Cascade-DNA complexes were introduced in the chamber with 500 μM ATP and 10 μM Co^2+^ while imaging at room temperature (23± 1 °C) and binding events were monitored in real time. For DNA unwinding assays the Cascade-DNA complexes were incubated for 5 minutes, allowing the complexes to form a stable interaction with the surface immobilized Cas3 molecules. Reeling was initiated by introducing ATP into the chamber while imaging at room temperature (23 ± 1 °C), allowing for visualization of the dynamics of Cas3 in real time. Notably, experiments were performed in absence of Co^2+^, unless explicitly stated in the text. To visualize the dynamics of Cas3, Cy3 molecules were excited an area of 50 × 50 μm^2^ with a 28% of the full laser power (9 mW) green laser (532nm), while the time resolution was set to 0.1 second. Under these imaging conditions we obtained a high signal-to-noise ratio that allowed us to visualize kinetic intermediates while imaging over time periods of 3.5 min. Under these conditions photobleaching of the donor and acceptor dye during our observation time was minimized.

### Single-molecule fluorescence data analysis

A series of CCD images were acquired with laboratory-made software at a time resolution of 0.1 sec. Fluorescence time traces were extracted with an algorithm written in IDL (ITT Visual Information Solutions) that picked fluorescence spots above a threshold with a defined Gaussian profile. The extracted time traces were analyzed using custom written MATLAB (MathWorks) and python algorithms. FRET efficiency was defined as the ratio between the acceptor intensity and the sum of the acceptor and donor intensities. The crosstalk between the two detection channels was not corrected to minimize any artefact in using the step-finder algorithm.

For dwell-time (Δ*τ*) analysis, the start and end of each reeling event was determined (Fig. 3a). The start of each event was marked by an abrupt decrease in the donor signal, whereas the end of each event was marked by an abrupt increase in the donor signal (Fig. 2b-c). Selecting the start and end of each event yielded the duration of each event, which was plotted in a histogram. These dwell-time distributions were fitted with a gamma distribution using maximum-likelihood estimations, which yielded an estimation of the number of hidden steps (N) and the rate per step (k). To obtain the global change in the velocity of Cas3 the number of hidden steps (N) and the rate per step (k) were converted to the effective rate (k^effective^, 1/Δ*τ*). The effective rate (k^effective^, 1/Δ*τ*) was obtained by dividing the rate per step (k) by the number of steps (N). Next, this effective rate was converted to velocity (bp/ s) by multiplying the effective rate by the FRET range of 22 base pairs (Extended Data Fig. 8a-b). The 95% confidence intervals (errors) of the dwell-times were obtained by empirical bootstrap analysis as described by^39^.

The step-size was characterized by adopting an automated step-finder algorithm, described previously by^28,34^. The step-finder algorithm yielded the average FRET value for each plateau, the size of each step in between the plateaus and the duration/ dwell-time (Δ*τ*) of each plateau. To be able to correlate the size of each step in FRET to distance in base pairs, a set of constructs was generated where the distance between donor and acceptor was systematically increased (Extended Data Fig. 8a-b). The slope of this calibration curve yielded a conversion factor, in which a change of ΔE=0.05 corresponds to a distance of one base pair. This allowed direct conversion of the step-size in FRET to distance in base pairs.

The dwell-time distributions for each step were fitted with a gamma distribution using maximum-likelihood estimations (MLE), which yielded an estimation of the number of hidden steps (N) and the rate per step (k). During MLE, each data point is weighted with equal importance. As a consequence, the minor populations in the tail of the distribution are given a substantial amount of priority during minimization of the fit. This causes the fit to widen, which results in an under-estimation of the number of steps and thereby an over-estimation of the rate per step. To correctly interpret data, only the data in the peak of the distribution was fitted, through the use a threshold (Extended Data Fig. 9a-g). Notably, the minor populations in the tail of the distribution may represent stalled helicases or enzymes that have a significantly slower velocity duo to static disorder^29^.

